# Cone-Opponent Ganglion Cells in the Primate Fovea Tuned to Non-Cardinal Color Directions

**DOI:** 10.1101/2023.09.15.557995

**Authors:** Tyler Godat, Kendall Kohout, Keith Parkins, Qiang Yang, Juliette E. McGregor, William H. Merigan, David R. Williams, Sara Patterson

## Abstract

A long-standing question in vision science is how the three cone photoreceptor types – long (L), medium (M) and short (S) wavelength sensitive – combine to generate our perception of color. Hue perception can be described along two opponent axes: red-green and blue-yellow. Psychophysical measurements of color appearance indicate that the cone inputs to the red-green and blue-yellow opponent axes are M vs. L+S and L vs. M+S, respectively. However, the “cardinal directions of color space” revealed by psychophysical measurements of color detection thresholds following adaptation are L vs. M and S vs. L+M. These cardinal directions match the most common cone-opponent retinal ganglion cells (RGCs) in the primate retina. Accordingly, the cone opponency necessary for color appearance is thought to be established in cortex. However, small populations with the appropriate M vs. L+S and L vs. M+S cone-opponency have been reported in large surveys of cone inputs to primate RGCs and their projections to the lateral geniculate nucleus (LGN), yet their existence continues to be debated. Resolving this long-standing open question is necessary because a complete account of the cone-opponency in the retinal output is critical for efforts to understand how downstream neural circuits process color. Here, we performed adaptive optics calcium imaging to longitudinally, noninvasively measure foveal RGC light responses in the living macaque eye. We confirm the presence of L vs. M+S and M vs. L+S neurons with non-cardinal cone-opponency and demonstrate that cone-opponent signals in the retinal output are more diverse than classically thought.

## Introduction

A goal of visual neuroscience is to determine how information is organized and represented at various stages of the visual system to produce our perception of the world. To understand color perception, a fundamental line of investigation focuses on determining how L-, M- and S-cones are combined at each stage. These questions are especially critical for retinal ganglion cells (RGCs), which provide the only visual input to the brain and thus make up the building blocks for all vision-related computations. As such, a full account of cone opponency in the retinal output is essential for a comprehensive understanding of downstream color processing.

In primates, the cones are combined to form three axes called the “cardinal directions of color space”. One direction responds to luminance (L+M) modulations, while the other two respond to isoluminant chromatic modulations (L vs. M and S vs. L+M). The cardinal directions were established by measuring detection thresholds following chromatic contrast adaptation (Krauskopf et al., 1982). A follow-up physiological survey found many LGN neurons are tuned to these three axes (Derrington et al., 1984) and established the widely-used DKL color space (**Figure 1A**; Brainard, 1996). In the retina, the cone inputs to the cardinal directions correspond to the three most common RGC types, the parasol, midget and small bistratified (Calkins, 2000). Based on this compelling results, the cardinal axes have served as the foundation for most color vision models (Lennie and D’Zmura, 1988; De Valois and De Valois, 1993; Stockman and Brainard, 2010).

**Figure 1.**
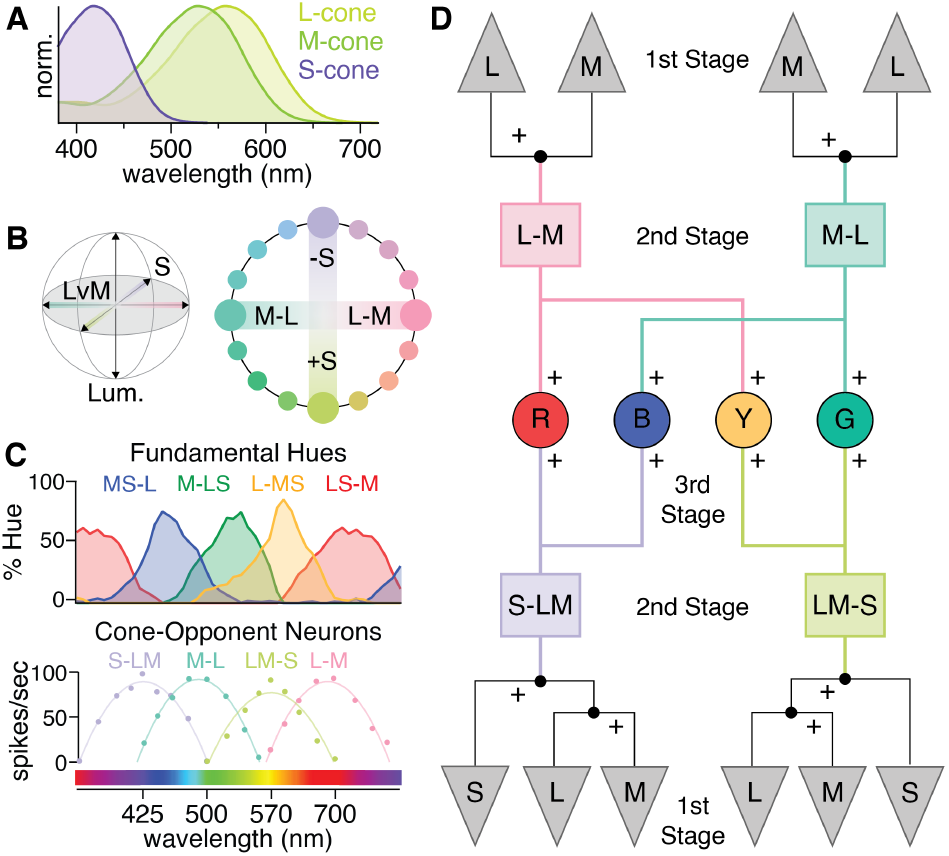
The most common cone-opponent neurons in the primate early visual system do not match hue perception. **(A)** Normalized spectral sensitivities for L-, Mand S-cones, which are are often called “red”, “green” and “blue”. However, each is individually colorblind as their outputs confound wavelength with intensity (Baylor et al., 1987). As a result, color vision requires comparing different cone types. **(B)** The isoluminant plane of the three-dimensional DKL color space (gray shading) formed by the cardinal L vs. M and S axes. Blue, yellow and green are not located along the cardinal axes and thus require both L vs. M and S-cone input. While red is near L–M in the DKL isoluminant plane, S-cone input is needed to explain the redness perceived in short-wavelength violet light. **(C)** Cone-opponent LGN neurons (bottom) initially linked L vs. M to “red-green” and S vs. LM to “blue-green” or “blue-yellow” (De Valois et al., 1966; Wiesel and Hubel, 1966). However, their responses and cone inputs do not align with the tuning curves for red, green, blue, and yellow obtained with hue scaling (top). Adapted from (De Valois, 2004). **(D)** One form of the multi-stage model, adapted from Stockman and Brainard, 2010. The cardinal directions reflect the second stage consistent with color detection and comparisons of their outputs form a third stage with cone opponency that can explain color appearance. Other multi-stage models omit the once-controversial S-OFF neurons and only use the output of S-ON small bistratified RGCs (De Valois and De Valois, 1993).

However, the cone inputs to the cardinal directions cannot explain color appearance (Stockman and Brainard, 2010) and do not match the cone opponency derived from psychophysical measurements of red-green and blue-yellow color appearance: L vs. M+S and M vs. L+S, respectively (Werner and Wooten, 1979; Shevell and Martin, 2017). The cardinal chromatic directions and corresponding cone-opponent neurons are often called “red-green” and “blue-yellow” (Lankheet et al., 1998a; Conway, 2001; Wool et al., 2018), but are perceived as red-cyan and lavender-lime (**Figure 1B**). While L–M appears red in the DKL isoluminant plane (Wuerger et al., 2005), the redness perceived in short-wavelength violet lights requires S-cone input (**Figure 1C**; Ingling, 1977; Fuld et al., 1981).

The currently-favored hypotheses propose cortical processing resolves the discrepancy between retinal cone opponency and color appearance (Lennie and D’Zmura, 1988; De Valois and De Valois, 1993; De Valois et al., 1997). In these “multi-stage” models (**Figure 1D**), the cones are the first stage, and L vs. M and S vs. L+M cone opponency form the second stage. A third stage accounts for color appearance with specific L vs. M and S-cone combinations.

An alternative hypothesis prompted by the confounded spectral and spatial information in L vs. M midget RGCs proposes that other RGCs with the cone opponency predicted by color appearance psychophysics contribute to downstream hue-encoding circuits (Rodieck, 1991; Calkins and Sterling, 1999; Schmidt et al., 2014; Neitz and Neitz, 2017). In these two-stage models, a greater diversity of second stage mechanisms replaces the third stage in **Figure 1D**. Most large-scale surveys have identified RGCs and LGN neurons with noncardinal L vs. M+S and M vs. L+S opponency (Padmos and Norren, 1975; De Monasterio, 1978; Malpeli and Schiller, 1978; Derrington et al., 1984; Valberg et al., 1986; Tailby et al., 2008; but see Sun et al., 2006). However, the existence of non-cardinal RGCs is not widely accepted and few investigations of L vs. M RGCs include S-cone stimuli. Regardless of their role in vision, confirming the existence of these RGCs would establish a greater diversity of cone opponency in the retinal output that challenges foundational assumptions of color vision models predicated on the existence of only two cardinal cone-opponent pathways.

The difficulty in confirming or denying the existence of RGCs with non-cardinal cone opponency stems from two technical challenges facing standard retinal physiology techniques. First, the fovea plays a major role in color vision, yet its fragility has largely limited *ex vivo* physiology to the periphery. Second, rarer primate RGC types are difficult to identify reliably in acute experiments. To overcome these challenges, we used adaptive optics and calcium imaging to non-invasively, longitudinally measure the responses of hundreds of individual foveal RGCs in the living macaque eye.

## Methods

### Animal care

Three macaques (*Macaca fascicularis*), two female and one male, were housed in pairs in an AAALAC accredited facility. All animals were in the care of the Department of Comparative Medicine veterinary staff of the University of Rochester’s Medical Center, including several full-time veterinarians, veterinary technicians, and an animal care staff who monitored animal health. Additional details on the vivarium environment are detailed in our previous work (McGregor et al., 2020; Godat et al., 2022). This study was carried out in strict accordance with the Association for Research in Vision and Ophthalmology (ARVO) Statement for the Use of Animals and the recommendations in the Guide for the Care and Use of Laboratory Animals from the National Institutes of Health. The animal protocol was approved by the University Committee on Animal Resources (UCAR) of the University of Rochester under PHS assurance number D16-00188 (A3292-01).

Each animal had no known history of ocular disease or vision abnormality. Each animal was fitted with a contact lens to enhance wavefront correction. The nasal retina of the left eye was imaged in M2 and the temporal retina of the right eye was imaged in M3 and M4.

The imaged eye of M2 had an axial length of 17.2 mm and dioptric power 57.5 D x 45°and 56.7 D x 135°. M3’s imaged eye had an axial length of 16.6 mm and dioptric power of 61.1 D x 85°and 61.8 D x 175°. M4’s imaged eye had an axial length of 16.9 mm and dioptric power of 60.1 D x 5°and 66.6 D x 95°.

### Viral delivery

All three macaques (M2, M3 and M4) received subcutaneous Cyclosporine A prior to injection of viral vectors. Blood trough levels were collected weekly to titrate the dosage to the range of 150–200 ng mL^−1^ and then maintained at that level. M2 began immune suppression in March 2018 with 6 mg kg^−1^, which was reduced to 4 mg kg^−1^ a month later. M3 started immune suppression in May 2019 at 6 mg kg^−1^, which was reduced a month later to 3.4 mg kg^−1^. M4 began immune suppression in December 2020 at 4 mg kg^−1^ and maintained this level.

The intravitreal injections used in this study have been described in full previously (Yin et al., 2011)) and are summarized below. The vector *AAV2-CAG-GCaMP6s* was synthesized by the University of Pennsylvania Vector Core. Injections were made into both eyes of each animal. Prior to injection, eyes were sterilized with 50% diluted Betadine, and the injection was made in the middle of the vitreous approximately 4 mm behind the limbus using a tuberculin syringe and 30-gauge needle. Following injection, each eye was imaged with a conventional scanning light ophthalmoscope (Heidelberg Spectralis) using the 488 nm autofluorescence modality to determine onset of GCaMP expression in the ganglion cell layer (GCL) and to monitor eye health. The data in this study was collected 1-year post-injection for M2, 1.3 years post-injection for M3 and 1.4 years postinjection for M4.

### Imaging preparation

Anesthesia and animal preparation procedures have been previously published (McGregor et al., 2020; Godat et al., 2022) and are briefly summarized here. The anesthesia and preparation were performed by a veterinary technician licensed by the State of New York (USA). All animals were fasted overnight prior to anesthesia induction the morning of an imaging session. During a session, animals were placed prone onto a custom stereotaxic cart and covered with a Bair Hugger warming system to maintain body temperature. Monitoring devices including rectal temperature probe, blood pressure cuff, electrocardiogram leads, capnograph, and a pulse oximeter, were used to track and record vital signs. Temperature, heart rate and rhythm, respirations and end tidal CO2, blood pressure, SPO2, and reflexes were monitored and recorded every fifteen minutes. Pupil dilation was accomplished using a combination of 1% Tropicamide and 2.5% Phenylephrine. A full list of all medications used and a full description of the anesthesia, intubation, pupil dilation, and recovery processes can be found in our previous work (McGregor et al., 2020).

### Adaptive optics scanning laser ophthalmoscopy

Data were collected using a fluorescence adaptive optics scanning light ophthalmoscope (AOSLO) previously described (Gray et al., 2006; Godat et al., 2022). The imaging paradigm is illustrated in **Figure 2**. Briefly, an 847 nm diode laser (QPhotonics) was used as a wavefront-sensing beacon with a Shack-Hartmann wavefront sensor to measure animals’ wavefront aberrations in real time (*∼* 20 Hz sample rate). A deformable mirror (ALPAO) with 97 actuators was used to correct the measured aberrations in closed loop at approximately 8 Hz across a 7.2 mm diameter pupil. A 796 nm center wavelength superluminescent diode (Superlum) was focused on the foveal cone photoreceptor mosaic to collect images (using a*∼* 2 Airy disk confocal image, 20 µm) used for navigating to the same retinal location in each experiment and for offline motion correction/registration of fluorescence images. Three LEDs (ThorLabs; center wavelengths and full-widths-at-half-maximum of 420 ± 7 nm, 530 ± 17 nm, 660 ± 10 nm) were presented to the foveal cones via a Maxwellian view stimulator (Westheimer, 1966; Baron, 1973). A 488 nm laser (Qioptiq) was focused on the GCL and used to excite fluorescence from GCaMP6-expressing cells, which was detected through a 520/35 nm filter (using a*∼* 2 Airy disk confocal pinhole, 20 µm in M2/M3, 25 µm in M4). The 488 nm excitation laser was presented only during forward raster scans of the AOSLO and only to the portion of the imaging field where RGCs were present (i.e., laterally displaced from the foveal cones and axially displaced to focus on the GCL). The somas of the foveal RGCs imaged lay at the margins of the foveal pit, approximately 1-4°from the foveal center. Using a previously published modeling approach (McGregor et al., 2018), these neurons are likely to be driven by cones within 36 arcmin eccentricity (within 120 µm radius from the foveal center).

**Figure 2.**
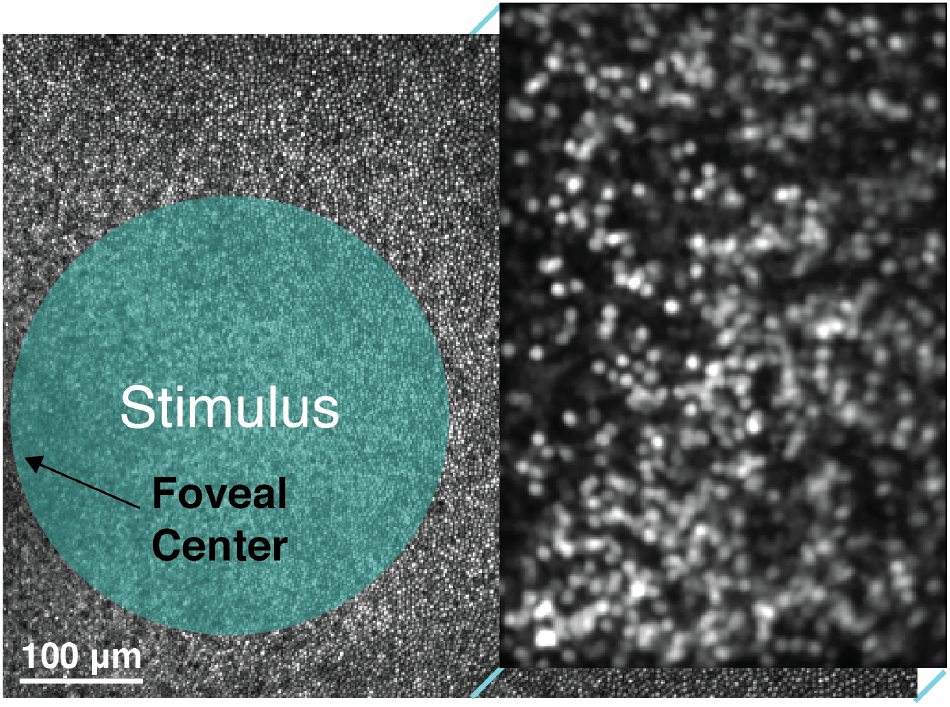
Experimental paradigm for stimulus delivery and calcium imaging in the living macaque fovea. Diagram illustrating the paradigm for in vivo calcium imaging with AOSLO. Data from M4 is shown with a 3.69 × 2.70° field of view. Reflectance imaging (796 nm) of the cone mosaic was performed across the full field of view while fluorescence imaging (488 nm) was restricted to the right half of the field of view and focused on GCaMP6s-expressing RGCs in the ganglion cell layer. Visual stimuli were focused on the foveal cones, which are displaced both laterally and axially from the RGCs. A wavefront sensing channel (847 nm) for real-time detection and correction of the eye’s monochromatic aberrations with adaptive optics is not pictured.

All light sources were delivered through a dilated pupil of*∼* 6.7 mm. The 488 nm excitation laser intensities on the retina were 3.2 µW/cm^2^ in M2, 1.7 µW/cm^2^ in M3, and 1.4 µW/cm^2^ in M4. Imaging fields of view subtended approximately 765 × 560 µm in M2, 740 × 540 µm in M3, and 740 × 540 µm in M4. The total retinal exposure to all light sources was calculated prior to each imaging session. Source powers measured at the animal’s pupil were nominally 7-15 µW at 488 nm, 250 µW at 796 nm, 30 µW at 847 nm and 5 µW from the three LEDs. The constant photopic adapting light presented with the LEDs mitigates residual responses to the spatially offset imaging light (Godat et al., 2022). Room lights were turned off during experiments to minimize ambient light.

The total exposure during each experiment was kept below the maximum permissible exposure for human retina according to the 2014 American National Standards Institute and most exposures were also below a reduced limit that was further scaled by the squared ra-tio of the numerical aperture of the human eye to the primate eye (*∼* 0.78-0.85). In this study, exposures during 2-4 hours of imaging ranged from 60-94% of the human retinal limit, with the majority between 60-85%. All sessions were spaced a minimum of 5 days apart per animal, so cumulative light exposure was not calculated.

### Visual stimuli

All stimuli were presented as a 1.3°(*∼* 250 µm) diameter spatially-uniform circle modulated sinusoidally in time at 0.2 Hz for M2 and 0.15 Hz for M3/M4. The low temporal frequency was chosen to accommodate the slow kinetics of GCaMP6s. Together, the temporal and spatial stimulus characteristics were likely ineffective for many non-midget RGC types with transient responses and/or strong surround suppression. These features as well as the stringent responsivity cutoff described below are major contributors to the relatively large number of nonresponsive cells (16.26%) in our dataset.

Each stimulus presentation was 90 seconds long, following a 30-second adaptation period. Stimulus presentations were separated by at least 20 seconds. The 488 nm laser was turned on at least three minutes prior to stimulus presentation and kept at a constant value throughout the experiment. Five stimuli were used in the study: an isoluminant L–M stimulus (only M2, 15% L-cone and 17% M-cone modulation), an L-cone isolating stimulus (used in M3 and M4 with 24 and 26% modulation, respectively), an M-cone isolating stimulus (used in M3 and M4 with 33% and 35% modulation, respectively), an S-cone isolating stimulus (used in all animals with 69%, 92% and 94% modulation in M2, M3 and M4, respectively) and an achromatic stimulus modulating all three cone types equally around the white point (all animals, 100% modulation). A control stimulus was also presented without modulation around the mean, which had the approximate chromaticity coordinates of (0.33, 0.33). Stimuli were calculated using the Psychophysics Toolbox (Brainard, 1997; RRID: SCR_002881) and calibrated as previously described (Godat et al., 2022). Within an imaging session, each stimulus was presented three times.

Data was collected from two experiments for M2 and M3 and three experiments for M4. Some aspects of the data from M2 and M3 were previously published in a larger study on spatial frequency tuning in midget RGCs (Godat et al., 2022). As described below, repeatability between imaging sessions was used in validating cone opponency, so the present study excludes a macaque with only one imaging session (M1) from the previously published study. The dataset from M4, qualitative technique for verifying optical isolation (**Figure 5**), calculations on S-cone RGC densities (**Figure 6**), and detailed investigation into the different cone opponent groups (**Figure 4**) are all novel to the present study.

### Data analysis

Fluorescence videos were co-registered with the simultaneously-acquired reflectance videos frame-by-frame with a strip-based cross-correlation algorithm (Yang et al., 2014). Registration was performed relative to a reference image of the cone mosaic, taken during the experiment and used for online stabilization. For each experiment, a registered fluorescence video from the control trial was summed across frames to create an image of the GCL for region of interest (ROI) segmentation using GIMP (RRID: SCR_003182). Unless otherwise stated, analyses below were performed in MATLAB (Mathworks, RRID: SCR_001622). Final figures were created in Igor Pro 9 (Wavemetrics, RRID: SCR_000325) and Illustrator (Adobe, RRID: SCR_010279). To analyze each sinusoidal stimulus, each ROI’s responses (average of three presentations per stimulus) were filtered using a Hann windowing function and then Fourier transformed using custom software (Frequency Analysis, available at https://osf.io/s9qw4. The phase and amplitude at the modulation frequency were used to quantify the strength and sign (ON-or OFF-dominant) of the response to each stimulus. Fourier amplitudes at the stimulus frequency were normalized for each cell by computing a signal-to-noise ratio (*SNR*):

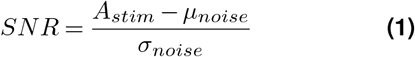

where *A*_*stim*_ is the Fourier amplitude at the stimulus frequency and *µ*_*noise*_ and *σ*_*noise*_ were the mean and standard deviation of the noise (0.32-1.08 Hz). The SNR response metric is equivalent to the sensitivity index d-prime (*d*^*′*^) that represents the response amplitude in SDs above noise (Horwitz, 2021). This approach enabled determination of an objective criteria for classifying responses as significant. A cutoff of two SDs above noise (dashed lines in **Figures 3** and **4**) was chosen for significance, which produced a 2.7% false positive rate across all ROIs in each experiment (i.e., 2.7% of all ROIs exceeded 2 SDs for the control stimulus). When each ROI’s responses were averaged across experiments, only one of the 486 cells imaged exceeded this cutoff for the control stimulus.

**Figure 3.**
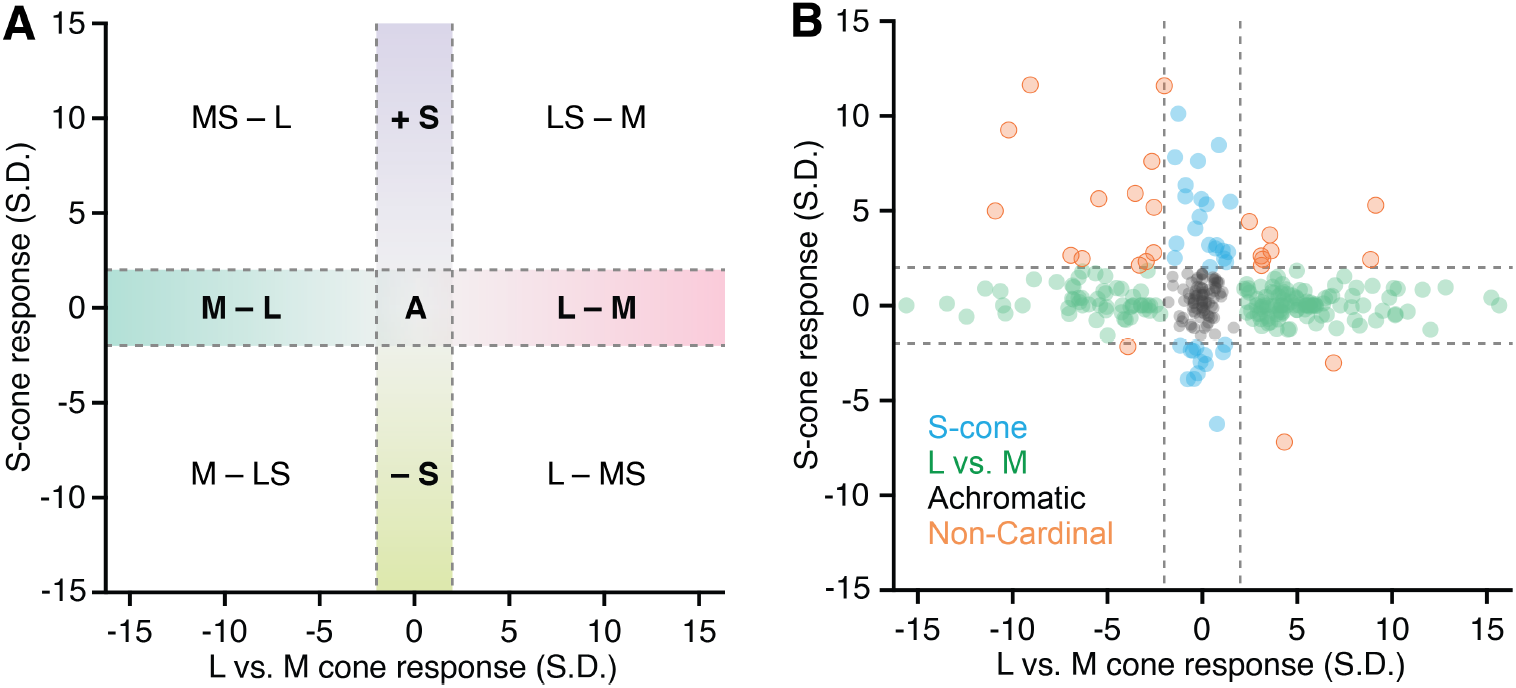
Functional classification of cone opponency in the living primate fovea. **(A)** The nine possible outcomes from classification criteria based on the sign and significance of L vs. M opponency and S-cone input. The axes correspond to the chromatic cardinal directions (L vs. M and S) and the dashed lines indicate the significance threshold of two standard deviations above noise. Neurons within the horizontal dashed lines have L vs. M opponency and lack significant S-cone input. Neurons within the vertical dashed lines have S-cone input but no significant L vs. M opponency. Neurons with both L vs. M opponency and S-cone input map to the four quadrants while neurons that only responded to the achromatic stimulus and not the cone-isolating stimuli fall in the center square labeled “A”. **(B)** 292 foveal neurons from three macaques classified based on their response at the stimulus frequency in units of standard deviations above noise (d-prime).

**Figure 4.**
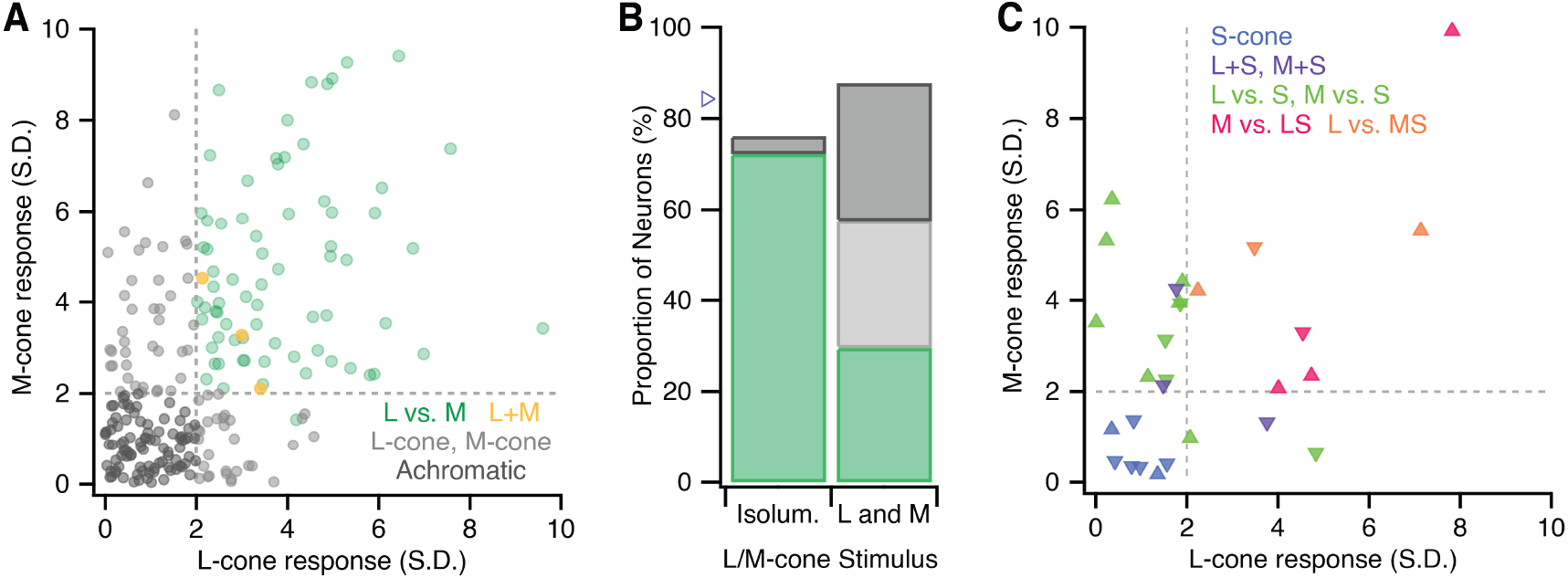
Detailed classification obtained with Land M-cone isolating stimuli in two macaques. Neurons with either Lor M-cone input as determined by the Land M-cone isolating stimuli used in M3 and M4 (254 of the 380 neurons analyzed). Note that the M-cone isolating stimulus had *∼*1.3x higher contrast than the L-cone isolating stimulus. **(A)** Neurons with only Lor M-cone input are shown in light gray. Achromatic and L vs. M opponent neurons from M3 and M4 shown in **Figure 3B** are replotted for reference. Due to the variability in Land M-cone inputs to midget RGCs, L vs. M midget RGCs could be located in any of the four sections (achromatic, L, M, or L vs. M). **(B)** The difference in of achromatic and L/M-cone response classification obtained with an isoluminant (L-M) stimulus (M2, n = 126) vs. Land M-cone isolating stimuli (M3/M4, n = 254). The blue triangle marks the estimated density of Land M-cone center midget RGCs (**Equation 7**). Colors are the same as in 4A. **(C)** The subset of S-ON and S-OFF neurons with either Lor M-cone input. The S-ON, S-OFF and L vs. M ± S neurons from M3 and M4 are replotted from **Figure 3B**. Triangles pointing upwards and downwards mark neurons with S-ON and S-OFF responses, respectively

**Figure 5.**
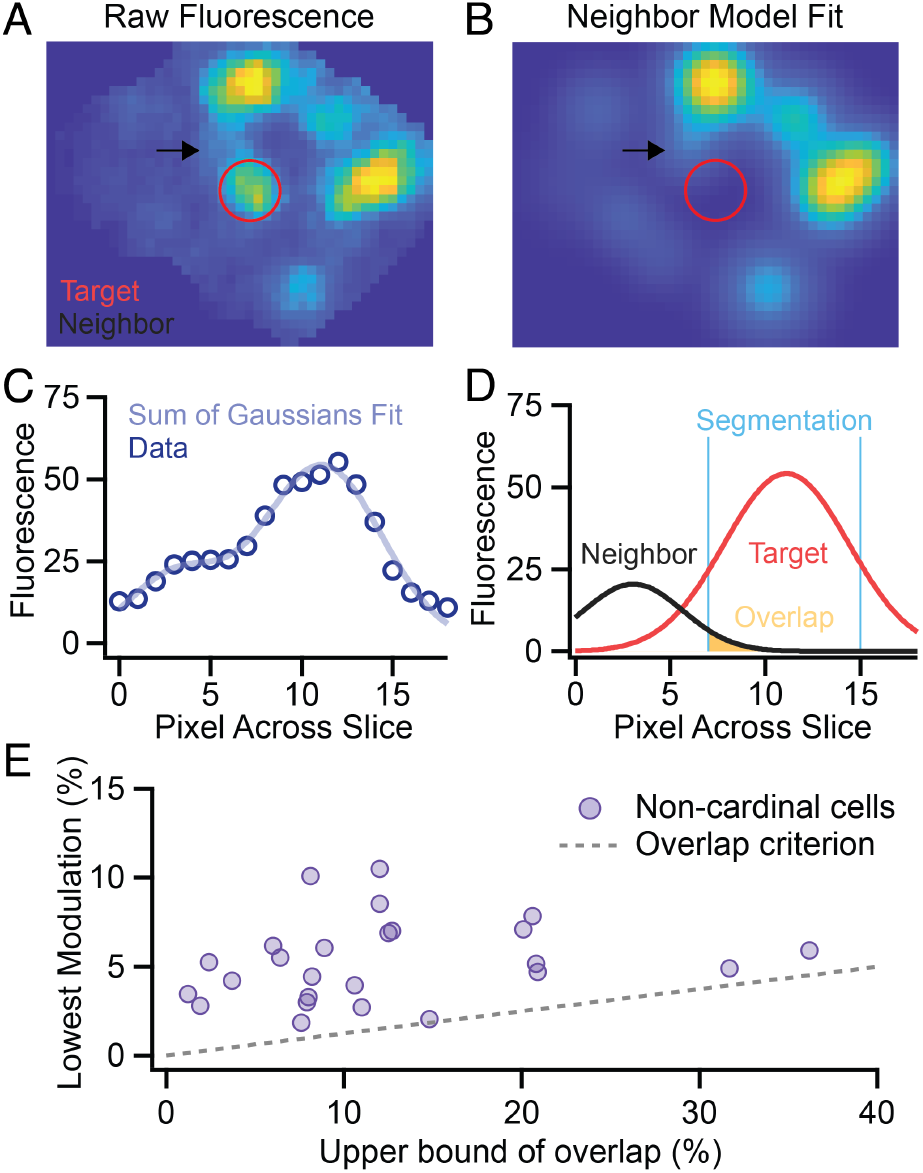
Sum of Gaussians model to verify optical isolation. **(A)** Fluorescence image of the neighborhood around an L vs. M ± S neuron (red). We chose this cell for demonstration due to the proximity of a neighbor cell (black arrow) and it is not representative of all L vs. M ± S neurons which typically have much less overlap. **(B)** The model fits a 2D Gaussian to each cell and the overlap is calculated from the area of neighboring cells within the target cell’s segmentation mask. The result is an upper bound on overlap because the 2D Gaussian fit overestimates the cell’s spatial footprint and the overlap from all neighboring cells was summed together. For display of overlap, the Gaussian representing the target cell is omitted. **(C, D)** A one-dimensional view of the fitting procedure and overlap calculation for the target cell and its nearest neighbor. **(E)** Comparison of the lowest modulation depth obtained across stimuli to the percent overlap from other cells within the target cell’s segmentation mask. The maximum modulation explainable by crosstalk was bounded by the product of the percent overlap and the largest modulation observed across experiments (gray dashed line).

**Figure 6.**
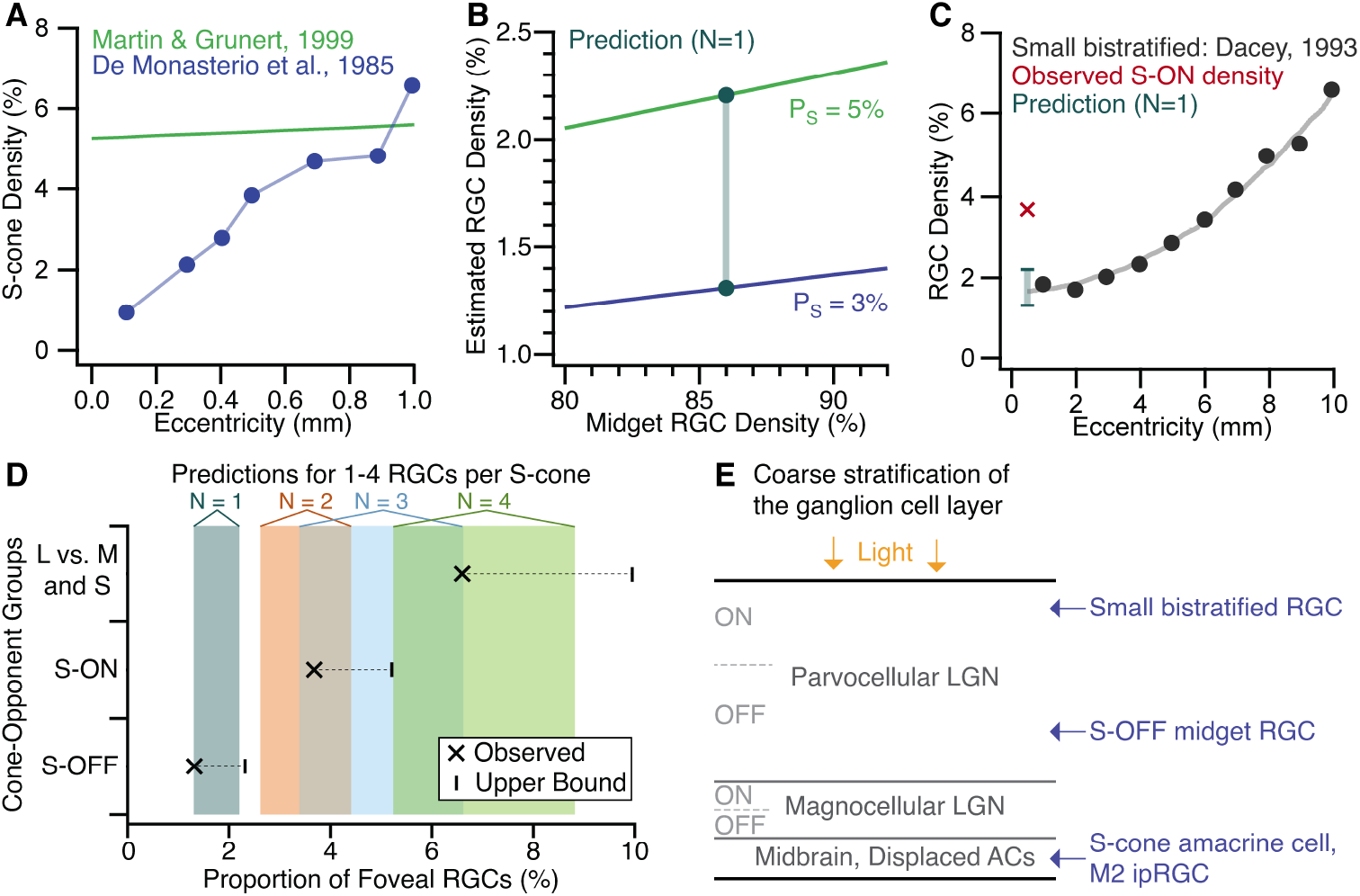
Modeling the density of foveal RGC types with S-cone input. Given the expected number of RGCs per S-cone (*N*), the model estimates that RGC type’s density (proportion of all foveal RGCs) from two parameters: S-cone density (*P*_*S*_) and midget RGC density (*p*_*mRGC*_ ; **Equations 5** and **6**). **(A)** Datasets used to determine the upper and lower bounds for S-cone density (De Monasterio et al., 1985; Martin and Grünert, 1999). **(B)** Density estimates at *N* = 1 for the upper and lower bounds on S-cone density over a range of midget RGC densities. The prediction range (teal) was obtained with *P*_*mRGC*_ = 0.86 (Peng et al., 2019). **(C)** Anatomical estimates of small bistratified RGC density (Dacey, 1993) fall within the prediction range from B, but the density of S-ON neurons from **Figure 3B** does not. **(D)** Estimated density ranges for 1-4 RGCs per S-cone (shaded boxes) compared with the observed densities of cardinal S-ON, cardinal S-OFF and L vs. M ± S neurons. The upper bounds for each group reflect the addition of L vs. S, M vs. S, M+S and L+S neurons (see main text). **(E)** Diagram (not to scale) of the foveal GCL’s stratification (Perry and Silveira, 1988) and known soma locations for S-cone neurons (Patterson et al., 2019a, 2020a, 2020b). The over-representation of cardinal S-ON neurons suggests the imaging plane was biased towards a more superficial layer of the GCL.

Between experiments, there were slight changes in axial focus, which can influence the measured strength of an ROI’s stimulus-driven response. However, these small focus changes could also increase and decrease the contributions of out-of-focus cells. Rather than try to distinguish between these alternatives, we chose to omit all cells without significant responses in at least two experiments (of two for M2/M3 and three for M4). This constraint likely leads to false rejections but increases confidence in those cells, which did respond significantly and reliably across experiments.

### Comparison of S and L vs. M response distributions

The two-sample Kolmogorov-Smirnov test was used to compare the distributions of L vs. M and S-cone responses in cardinal and non-cardinal cone-opponent neurons. The null hypothesis evaluated was that the responses of the non-cardinal cells come from the same underlying distribution as the cardinal cells. Because the L vs. M and S response distributions were discontinuous due to the absence of reliable data between -2 and 2 SDs, L–M, M–L, S-ON and S-OFF responses were considered separately.

The following criterion was used to determine whether the sample sizes were sufficient for reliable use of the two-sample Kolmogorov-Smirnov test:

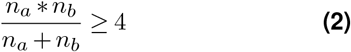

where *n*_*a*_ and *n*_*b*_ are the number of samples in the two distributions. The neurons with S-OFF responses were not evaluated because they failed to meet this criterion. For L–M and M–L, responses of neurons with both L vs. M opponency and S-cone input were compared to the distribution of responses from neurons with L vs. M opponency that lacked significant S-cone input. For S-ON responses, neurons with both S-cone input and L vs. M opponency were compared to cardinal S-cone neurons lacking significant L vs. M opponency.

### Sum of Gaussians model

To calculate the upper bound for overlapping light distributions contributing responses to a single segmented ROI, we developed a sum of Gaussians model. This model allowed quantification of the possibility that cells with non-cardinal cone opponency reflect the responses from two more common cone-opponent neurons, one with S-cone responses and one with L vs. M cone opponency. For each target cell, a new mask was manually placed centered on the target cell and including the nearest 8-14 neighboring cells (many of which were not initially segmented due to evidence for poor optical isolation in at least one experiment). This mask was applied to the fluorescence image to restrict fitting to the neighborhood around each target cell.

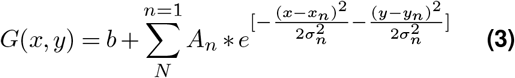

where *b* is the minimum background pixel value within the mask, *N* is the number of ROIs in the image, *A*_*n*_ is the Gaussian amplitude for the *n*-th cell, (*x*_*n*_, *y*_*n*_) is the center coordinates of the *n*-th Gaussian, and *a*_*n*_ is the standard deviation of the *n*-th Gaussian. A nonlinear least squares fit was used to match the sum of Gaussians model to the measured data. For the fit, (*x*_*n*_, *y*_*n*_) were fixed while *A*_*n*_ and *a*_*n*_ were free parameters. This method was chosen because the center of each cell was clearly identifiable and allowing (*x*_*n*_,*y*_*n*_) to vary freely would have required an arbitrarily high number of Gaussians for a given image. Contrast enhancement was employed during cell identification to ensure all cells were identified and assigned x and y values. A lower bound of unity was set for both *A*_*n*_ and *a*_*n*_. The images ranged from 0-255, so a Gaussian could not have *A*_*n*_ *<* 1 and *a*_*n*_ *<* 1 (soma size less than *∼* 1.5 µm) is inconsistent with published reports of foveal RGC somas (Curcio and Allen, 1990). Pixels outside the mask, but within the mask’s bounding box were set to the minimum value within the mask (*b*) to assist with fitting (**Figure 5A**).

Once a fit was calculated, the original segmentation mask of the target cell was applied to the modeled fit, and the overlap percentage was calculated as the sum of pixel intensities from nearest neighbors within the segmentation divided by the total sum of pixel intensities within the ROI (**Figure 5B-D**). This value, expressed as a percentage, gives an upper bound on how much light could have fallen within the ROI that originated from neighboring cells. This value represents an upper bound because cells have definite sharp borders, and despite a modest amount of optical blur and scattered light, their light distributions lack the spatial extent described by a Gaussian function. Thus, the overlap metric is a conservative estimate of potential optical crosstalk.

To interpret the upper bounds of overlap for each L vs. M ± S cell, we compared the overlap to the modulation depth for each stimulus (**Figure 5E**). The Fourier amplitude at the frequency stimulation and the DC offset (Fourier amplitude at the zero frequency) were obtained from the one-sided power spectrum and used to compute the modulation depth (*M*_*s*_) for each stimulus (*s*) in the following way:

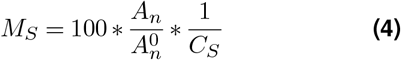

where *A*_*n*_ is the Fourier amplitude of the *n*-th cell at the modulation frequency, 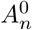 is the DC offset of the *n*-th cell, and *C*_*S*_ is the nominal cone modulation of stimuluss. In M2, *C*_*L−M*_ was 0.153 and *C*_*S*_ was 0.688. In M3, *C*_*L*_ was 0.243, *C*_*M*_ was 0.325 and *C*_*S*_ was 0.92. In M4, *C*_*L*_ was 0.258, *C*_*M*_ was 0.354 and *C*_*S*_ was 0.939.

The maximum modulation explainable by optical crosstalk for each L vs. M ± S cell is the product of the percent overlap and the maximum modulation across all segmented neurons (dashed line in **Figure 5E**). If the cell’s lowest modulation depth exceeded this value, then their responses could not be explained by optical crosstalk from neighboring cells.

### Density estimates for S-cone RGCs

To provide context for interpreting the densities of the cardinal and non-cardinal neurons obtained in our study, we developed an approach for estimating the density of an RGC type based on their density relative to the Scones. In other words, the model estimates the density of an RGC type given the expected number per S-cone (*N*) and is agnostic to the underlying S-cone circuitry.

For example, even though small bistratified RGCs receive input from multiple S-cones, there is roughly one small bistratified RGC per S-cone (*N* = 1) (Calkins et al., 1998; Schein, 2004). Given a measured or predicted value for *N*, the density of an RGC type (*D*_*S*_) can be calculated as follows:

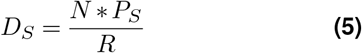

where *R* is the ratio of RGCs to foveal cones and *P*_*S*_ is the fraction of foveal cones that are S-cones. Thus, the calculation depends critically on two values from the literature: *P*_*S*_ and *R*. The most challenging parameter is the RGC-cone ratio, which varies within and between individuals and is complicated by the displacement of RGCs from their cone inputs (Curcio and Allen, 1990). Moreover, published values for macaque foveal RGC-cone ratios are highly variable, ranging from 2.15:1 to over 3:1 (Schein, 1988; Wässle et al., 1998). We instead estimated the RGC-cone density by relying on the well–established numbers of midget RGCs per foveal cone: 2 per L/M-cone and 1 per S-cone (Calkins et al., 1994; Patterson et al., 2019a). This stereotyped circuitry can be used to estimate the number of RGCs per foveal cone as:

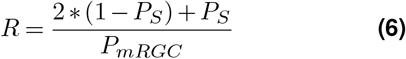

where *P*_*S*_ is the fraction of all cones that are S-cones and *P*_*mRGC*_ is the fraction of all foveal ganglion cells that are midget RGCs. The estimates in the text used *P*_*S*_ = [0.03, 0.05] and *P*_*mRGC*_ = 0.86. The macaque foveal S-cone density range was obtained from a digitized figure (Calkins, 2001). The lower bound of 3% reflects the average S-cone density from 0-500 µm in (De Monasterio et al., 1985). The upper bound of 5% reflects the average density over the same eccentricities for the temporal and nasal datasets (Martin and Grünert, 1999). The foveal midget RGC density (86%) comes from transcriptomics analysis of multiple macaque foveas and thus is more robust to variation within and between individuals (Peng et al., 2019). Digitized values were rounded to the nearest percentage due to the limited resolution of the figures. Confidence in this approach is provided by the strong agreement between the RGC-cone ratios obtained with **Equation 6** (2.26:1-2.29:1 for an S-cone density range of 3-5%) and recent empirical reports of 2.2:1 in the human fovea (Masri et al., 2020).

A similar approach was used to estimate the proportion of macaque foveal RGCs that are L/M-cone center midget RGCs (*D*_*LM*_):

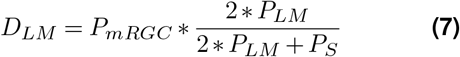

where *P*_*LM*_ = 1*− P*_*S*_ and represents the fraction of all cones that are Lor M-cones. Unlike the S-cone RGC density estimate above, this calculation did not require determination of the RGC-cone ratio and, as a result, was less dependent on the value of *P*_*S*_.

Importantly, these calculations depend on the stereotyped “private line” circuitry of midget RGCs (Kolb, 1970; Kolb and Marshak, 2003) and are only valid for the fovea and central retina. The code used for the density estimates is publicly available at https://github.com/sarastokes/NonCardinalRGCs.

## Results

### Longitudinal imaging of foveal ganglion cell physiology in the living macaque eye

We leveraged a powerful in vivo retinal physiology technique that combines adaptive optics scanning laser ophthalmoscopy (AOSLO) with calcium imaging to present visual stimuli and image RGC responses at the center of the living macaque fovea (Gray et al., 2006; Yin et al., 2014; McGregor et al., 2018; Roorda et al., 2022). GCaMP6s expression in the foveal ganglion cell layer was obtained through intravitreal injections of *AAV2:CAG:GCaMP6s* (Yin et al., 2011; Chen et al., 2013). The imaging paradigm is depicted in **Figure 2** and described in full in the **Methods**. Briefly, the cellular resolution necessary to non-invasively image calcium responses in vivo was achieved with adaptive optics, which detects and corrects for the eye’s monochromatic aberrations in real time (Liang et al., 1997; Williams et al., 2023). Reflectance imaging of the cone mosaic was used for online eye tracking, offline registration, and navigation to the same retinal location across imaging sessions. Fluorescence imaging of GCaMP6sexpressing foveal RGCs was performed while spatially uniform chromatic visual stimuli were presented to the cones with a Maxwellian view stimulator. Due to the natural spatial displacement of foveal RGCs from their cone inputs, the imaging light was displaced both laterally and axially from the RGCs’ cone inputs and the visual stimulus.

The unique advantages of this approach are well suited to resolving the question of whether neurons tuned to non-cardinal color directions are present in the primate retina. The fovea plays a key role in color perception, yet its fragility poses a major obstacle for *ex vivo* electrophysiology. As a result, our understanding of foveal cone-opponent RGCs is largely inferred from anatomy and their response properties in the retinal periphery, despite evidence that the physiology of foveal neurons often differ from their peripheral counterparts in unexpected ways (McMahon et al., 2000; Sinha et al., 2017; Lee et al., 2018; Baudin et al., 2019). With our noninvasive calcium imaging approach, we were able to study foveal RGCs in their natural habitat (the living eye). Simultaneously recording from hundreds of foveal RGCs increased the likelihood of finding rarer cone-opponent RGC types. Moreover, we can return to the same RGCs in subsequent sessions to demonstrate the reliability of our measurements and study the rarer types in detail. These advantages enabled us to provide new insight into the classic question of cone-opponency in the primate retinal output.

### Classification of cone opponency in foveal neurons with *in vivo* calcium imaging

In total, we analyzed 486 well-isolated neurons across three macaques (M2, M3 and M4). The classification was based on responses to achromatic, isoluminant and cone-isolating stimuli modulated sinusoidally in time. The amplitude and phase at the modulation frequency were used to quantify the strength and sign (ON-or OFF-dominant) of the L-, M- and S-cones input to each neuron. We normalized the response amplitudes obtained in each session by calculating a signalto-noise (SNR) metric equivalent to d-prime, which quantitatively represents the strength of the neuron’s calcium response relative to noise. A key advantage of the SNR metric was that it enabled objective determination of whether a neuron responded significantly to each stimulus. A highly stringent cutoff of two standard deviations (SDs) above noise was established using control data (see **Methods**). Using this criterion, 83.7% (407/486) of the neurons responded significantly to at least one stimulus when averaged across experiments. 78.2% (380/486) responded reliably between experiments, which were separated by a week or more. The 106 neurons that either did not respond significantly to any stimulus or exhibited unreliable responses to one or more stimuli were omitted from all subsequent analyses.

We classified the 380 reliably responsive foveal neurons based on three criteria: 1) the presence of significant L vs. M opponency, 2) the presence of significant S-cone input, and 3) the sign (ON- or OFF-dominant) of each significant input. The nine potential outcomes are plotted on the cardinal axes as illustrated in **Figure 3A**, where the gray dashed lines indicate the significance cutoff of ±2 SDs above noise. Together, the vertical and horizontal dashed lines in **Figure 3A** mark the boundaries of the cardinal axes. Neurons with L- and M-cone input of opposite signs (L vs. M opponency) and no S-cone input fall within the horizontal dashed lines while neurons with S-cone input that lack L vs. M opponency fall within the vertical dashed lines. Neurons with non-cardinal cone opponency map onto one of the four quadrants, depending on the sign of each cone input, and together can be classified as having non-cardinal cone opponency.

For two of the three macaques studied, L- and M-cone isolating stimuli were used instead of an L-M isoluminant stimulus. The L/M-cone isolating stimuli were adopted to further increase confidence in identified non-cardinal neurons. These stimuli revealed 84 neurons with significant responses to either L-or M-cone isolating stimuli, 14 of which had S-cone input. Together, these neurons made up 33.5% of the 274 neurons tested with the L- and M-cone isolating stimuli and 22.4% of the full dataset of 380 neurons. These L-or M-cone neurons could not be accurately plotted along the L vs. M axis in **Figure 3** and are instead plotted on L- and M-cone axes (**Figure 4**).

### Heterogeneous L- and M-cone input to foveal midget RGCs

L vs. M neurons were the dominant cone-opponent population in our dataset, making up 43.7% (166/380) of the classified neurons. In the 274 neurons tested with L- and M-cone isolating stimuli, 30 responded to the L-cone isolating stimulus only and 41 to the M-cone isolating stimulus only (**Figure 4A**). Together, these L-or M-cone neurons made up 18.7% of the 380 neurons analyzed. There is no direct evidence for retinal circuits that distinguish between L-or M-cones to contact just one of the two types; however, the L- and M-cone center midget RGC (L/M midget RGCs) receptive fields can be biased towards one type. In the fovea, their center-surround receptive fields compare the photon catch in the single L-or M-cone center to the photon catch in neighboring L/M-cones in the surround (reviewed in Patterson et al., 2019b). As the relative numbers of L- and M-cones in the surround receptive field are largely random, L-cone center midget RGCs, for example, can vary from strongly cone-opponent (only M-cones in the surround) to achromatic (only L-cones in the surround). Given the strict significance cutoff employed in our classification, an M-cone center RGC with few L-cones in the surround receptive field would respond significantly to the isoluminant and M-cone isolating stimuli, but not the L-cone isolating stimulus (or vice versa for an L-cone center RGC with little to no spectral opponency).

The possibility that the L-or M-cone neurons are midget RGCs with minimal spectral opponency is supported by the difference in L vs. M neurons detected with the isoluminant stimulus (72.2% in M2) compared to the L- and M-cone isolating stimuli (30.3% in M3 and M4; **Figure 4B**). Collectively, the L vs. M, L-cone and M-cone neurons in M3 and M4 made up 57.5% of the neurons analyzed, which is more comparable with the frequency of L vs. M neurons obtained in M2. However, these densities are lower than the estimated proportion of foveal RGCs that are L-or M-cone center midget RGCs, which is *∼*84% (**Equation 7**).

Interestingly, only three neurons had significant re-sponses to both the L- and M-cone isolating stimulus that were in phase (L+M). All three had ON responses (**Figure 4A**). The largest population of L+M neurons, ON and OFF parasol RGCs, are estimated at*∼* 6% of the foveal output (Ma et al., 2023). This discrepancy is likely due to stimulus choice. To accommodate population-level data collection and Fourier analysis with a slow calcium indicator, we used a large spatially uniform stimulus (1.3°diameter) and low tem-poral frequencies (0.15–0.2 Hz). Together, these features made our stimuli ineffective for neurons with transient responses and/or strong surround receptive fields, which are common features of parasol RGCs and other magnocellular LGN-projecting RGCs (Derrington and Lennie, 1984; Solomon et al., 2006).

Nonetheless, 82 (21.6%) neurons only responded significantly to the achromatic stimulus (center of **Figure 3A**) and bottom left of **Figure 4A**). While these neurons could include parasol RGCs, their high density suggests many are instead midget RGCs (**Figure 4B**). The contrast of the achromatic stimulus was substantially higher than the isoluminant, L-cone and M-cone isolating stimuli. As a result, midget RGCs with weak spectral opponency may respond well to the high contrast achromatic stimulus, but lack sufficient L vs. M opponency to generate significant responses to the lower contrast isoluminant and L/M-cone isolating stimuli. Collectively, the densities of achromatic, L vs. M, L-cone and M-cone neurons (83.95%) was consistent with the estimated *∼* 84% density of L- and M-cone center midget RGCs in the macaque fovea.

### Diverse S-cone opponency in foveal neurons

We identified a smaller population of neurons falling along the cardinal S-cone axis: 14 (3.7%) with S-ON responses and 5 (1.3%) with S-OFF responses. The density of S-ON neurons is higher than expected of small bistratified RGCs (**Figure 6C**), the most common foveal S-ON neuron. This discrepancy could reflect the location of the imaging plane within the GCL (discussed below) or the inclusion of rarer S-ON neurons, such as large bistratified RGCs, ON narrow thorny RGCs, M2 intrinsically photosensitive RGCs and displaced S-cone amacrine cells (Dacey and Packer, 2003; Patterson et al., 2020b, 2020a; Mazzaferri et al., 2023).

Of the 380 neurons studied, 25 (6.6%) exhibited both L vs. M opponency and S-cone input and thus were not located along either cardinal axis: 13 MS–L, 8 LS–M, 3 L–MS, and 1 M–LS. Because their role in vision remains unclear, we refer to the four groups collectively as L vs. M ± S neurons. The signed cone inputs to these four groups are consistent with the signed cone inputs to red (LS–M), green (M–LS), blue (MS–L), and yellow (L–MS) derived from hue scaling experiments in **Figure 1C**. Importantly, this correlation does not imply causation. The relative frequencies of the four cone-opponent groups differed with the two groups receiving S-ON input (LS-M and MS-L) outnumbering the two groups with S-OFF input (L-MS and M-LS).

In M3 and M4, the use of L- and M-cone isolating stimuli enabled identification of 14 (3.7%) S-cone neurons with significant responses to either the L-or M-cone isolating stimulus (6 S-ON and 8 S-OFF; **Figure 4C**). It is notable that over half of these neurons had S-OFF input, given the low numbers of cardinal and non-cardinal S-OFF responses in **Figure 3B**. Like the L-or M-cone neurons in **Figure 4A**, these neurons may have weak responses from the missing L-or M-cone that eluded detection. The interpretation of this population depends on whether the S-cone input is in phase with the L-or M-cone input.

Eleven (2.9%) neurons had S-cone input that was opponent to L-or M-cone input (**Figure 4C**). Nine neurons had M vs. S responses (6 S-ON/M-OFF and 3 M-ON/S-OFF) and two neurons had L vs. S responses (both L-ON/S-OFF). Given the reduced detection of L vs. M neurons in the two macaques where L- and M-cone isolating stimuli were used (**Figure 4B**), it is possible many of these neurons would have responded significantly to the more effective isoluminant stimulus. However, the L vs. S and M vs. S neurons could also be cardinal S vs. (L+M) neurons like the small bistratified and S-OFF midget RGC. Because these two possibilities could not be distinguished, the L vs. S and M vs. S neurons cannot be classified as “cardinal” or “non-cardinal”.

Another three neurons had M+S or L+S input: 2 MS-OFF and 1 LS-OFF (Figure 4C). These neurons are distinct from the cardinal S vs. L+M cells because their S-cone response is in phase with L or M. Including the M+S and L+S neurons, the total density of neurons tuned to non-cardinal color directions was 7.4% (28/380).

While the 14 S-cone neurons with either L-or M-cone input were not classified as S vs. L+M or L vs. M ± S, they can be used to establish the upper bounds of the underlying densities of cardinal and non-cardinal S-cone RGCs (triangles in **Figure 6D**). If the M+S, L+S, S vs. L, and S vs. M neurons in **Figure 4B** have a similar interpretation to the neurons with either L-or M-cone input in **Figure 4A** (weak L vs. M opponency), then total density of non-cardinal neurons could be as high as 10% (38/380; 14 MS-L, 14 LS-M, 5 M-LS, 5 L-MS). If the L vs. S and M vs. S neurons are part of the cardinal axis, the density of S-ON and S-OFF neurons could be as high as 5.3% and 2.4%, respectively. The S-ON and S-OFF cardinal neurons are considered separately because they correspond to distinct cell types (Thoreson and Dacey, 2019). Because the cell type(s) corresponding to the L vs. M ± S neurons are unknown, it is not clear whether or how this population should be divided. Taken together, these results confirm the presence of L vs. M ± S neurons in the primate fovea. Because they have both S-cone input and L vs. M cone-opponency, these neurons are distinct from the well-studied S vs. L+M and L vs. M RGCs and are not located along the cardinal axes. These data speak to the predictions two common color vision models make regarding cone opponency in the retinal output. As described in the Introduction, L vs. M ± S RGCs are central to one hypothesis (Schmidt et al., 2014) while the multi-stage models (e.g., **Figure 1D**) emphasize only three independent, orthogonal combinations of the L-, M- and S-cones (L vs. M, S vs. L+M and L+M) in the retinal output and propose L vs. M ± S opponency arises downstream (De Valois and De Valois, 1993; Stockman and Brainard, 2010). Importantly, this experiment did not test the predictions each model makes about how the different cone-opponent neurons contribute to color appearance, which will require the development of new techniques capable of performing causal manipulations that directly link cone-opponent RGCs to downstream visual functions or perception.

### Confirming the optical isolation of neurons with L vs. M opponency and S-cone input

A common concern with functional imaging is the possibility of response contamination from nearby neurons. Could the neurons identified with L vs. M ± S opponency reflect the responses of two overlapping neurons, one with L vs. M opponency and one with Scone input? Here we manually segmented only cells that were laterally well-isolated across all imaging sessions, as previously described (Godat et al., 2022). Any out-of-focus cell eluding detection during segmentation would be very dim and expected to contribute weakly to a segmented cell’s response. Our stringent responsivity criterion was chosen to eliminate such weak responses. Less than half of all visible cells in each region imaged met our criteria for segmentation and analysis. While these approaches have been successful in our past work, we sought to develop an additional method for quantitative verification of optical isolation.

We quantified optical isolation with a sum of Gaussians model for each L vs. M ± S neuron and its nearest neighbors (**Equation 3**). The model estimates an upper bound on the amount of fluorescence within each cell’s segmentation mask that could have originated from neighboring cells. We fit each target L vs. M ± S neuron and their nearest neighbors with twodimensional Gaussians (**Figure 5A-C**), then calculated the total overlap from all neighboring cells within the target cell’s segmentation mask (**Figure 5D**). The overlap was compared to the modulation depth of the tar-get cell’s weakest response to the cone-isolating stimuli (**Figure 5E**). The maximum response that could be explained by crosstalk was determined as the product of the percent overlap and the largest modulation depth recorded across all neurons in the dataset. The response amplitudes for all 25 L vs. M ± S neurons exceeded this threshold (dashed line in **Figure 5E**), indicating that their responses cannot be explained by crosstalk from nearby cells.

This model defines a highly stringent criterion that overestimates crosstalk in several ways. First, fitting somas with a two-dimensional Gaussian ignores the sharper borders of each cell and overestimates the spatial extent of each cell’s light distribution, thus providing a generous estimate of the amount of overlap between cells that might have occurred. Second, the overlap percentage for each cell reflects the summed contributions of all neighboring cells. The stimulus responses of each neighboring cell were not considered, as many were not segmented due to insufficient optical isolation (e.g., the neurons in the top right corner of **Figure 5A**). Many neighboring cells could have been omitted based on their response properties (e.g., a neighbor that did not respond to the S-cone isolating stimulus is unlikely to contribute to the target cell’s S-cone response), which would have reduced the summed overlap percentages plotted in **Figure 5E**. However, because all 25 L vs. M ± S cells passed our initial overly conservative test, these factors were not considered further.

### Variability within and between cone-opponent groups

Our experimental design focused on identifying statistically significant cone inputs, which naturally led to a discrete classification based on the presence or absence of L vs. M opponency and S-cone input. However, the boundaries in **Figure 3A** do not necessarily correspond to specific cell types. Indeed, L/M midget RGCs were likely distributed across the middle three sections of **Figure 3A** (L-M, M-L, achromatic) and across all four sections of **Figure 4A** (achromatic, L, M, L vs. M). The representation in **Figure 4A** is more comprehensive and consistent with the largely random arrangement of Land M-cones in midget RGC surround receptive fields (Paulus and Kroger-Paulus, 1983; Wässle et al., 1989; Lennie et al., 1991; Jusuf et al., 2006). An important direction for future research will be to determine whether the L vs. M ± S neurons are discrete groups or the tails of a distribution of S-cone input to L vs. M neurons.

We also observed variability within many of the cone-opponent groups. In interpreting **Figure 3B**, it is important to recognize that the cardinal axes were designed to facilitate comparison with the properties of psychophysically-identified second-stage mechanisms rather than to provide a precise representation of L-, Mand S-cone weights (Lankheet et al., 1998b). The representation in **Figure 3B** over-emphasizes the variability of L vs. M ± S neurons while de-emphasizing the variability of L vs. M and S vs. L+M neurons. Specifically, non-cardinal neurons appear more widely distributed than the cardinal neurons because their responses vary significantly along two dimensions rather than one. Indeed, a similar distribution is observed for the cardinal L vs. M neurons when their L- and M-cone responses are not condensed into a single dimension (top right section in **Figure 4A**).

To quantify these observations, we asked whether the distributions of L vs. M and S responses in non-cardinal cells were consistent with the distributions of L vs. M and S responses along the cardinal axes. The twosample Kolmogorov-Smirnov test failed to reject the null hypothesis that the L–M, M–L and S-ON responses of the non-cardinal and cardinal cells arose from different underlying distributions: L–M (p = 0.275), M–L (p = 0.209) and S-ON (p = 0.204). The small sample sizes of cardinal and non-cardinal S-OFF neurons prohibited reliable evaluation (**Equation 2**). The comparable distributions for S-ON and L vs. M responses could indicate shared underlying mechanisms (random wiring for L vs. M and selective wiring for S-ON). Indeed, the response distributions for L vs. M neurons and S-ON neurons were significantly different (p = 0.001). However, this important control is qualified by a potential confound: the contrast of the S-cone isolating stimulus was much higher than the contrasts of the L/M-cone isolating and isoluminant stimuli.

Taken together, these analyses indicate that the variability along each cardinal axis is similar for both the cardinal and non-cardinal neurons. As a result, combining the outputs of the cardinal RGCs to form cortical L vs. M ± S neurons, as proposed by multi-stage models (e.g., **Figure 1D**), would yield a similarly broad range of responses as is observed for the L vs. M ± S neurons in **Figure 3B**. More generally, many color vision models implicitly assume homogenous L-, Mand Scone weights for each individual neuron within a cone-opponent group. These models may benefit from incorporating more of the underlying heterogeneity in L-, M- and S-cone weights as efforts to incorporate variable tuning in other areas of visual neuroscience has produced more accurate and nuanced models (Cohen and Zaidi, 2007; Rokem and Silver, 2009). Psychophysical experiments such as the hue scaling in **Figure 1C** engage thousands of RGCs and the weights obtained could reflect the average of a cone-opponent population with variable L-, M- and S-cone weights. Determining whether or how the variability in cardinal and non-cardinal cone opponency influences color perception will be an important direction for future investigation.

### Assessing the densities of S-cone RGCs obtained with *in vivo* calcium imaging

A challenge in studying rarer neurons is determining whether the small sample sizes obtained in any one study are truly representative of the underlying population. This question is particularly relevant for relatively new techniques like *in vivo* calcium imaging where potential limitations may not be fully understood. It is worth reviewing three caveats to the densities obtained in our classification. First, we chose a highly stringent significance cutoff to maximize confidence in identified cone-opponent neurons rather than detection of all cone-opponent neurons. Second, the use of L- and M-cone isolating stimuli in two of the three macaques resulted in reduced sensitivity to L vs. M opponency and lower densities of both cardinal and non-cardinal L vs. M neurons (**Figure 4**). Third, the location of the imaging plane can introduce sampling biases as somas within the foveal GCL are coarsely stratified (Perry and Silveira, 1988). Because the imaging plane intersects the foveal slope, which begins as a monolayer and gradually increases to *∼* 6 cells deep, neurons from all areas within the GCL are imaged. As a result, sampling biases are mitigated, but not fully prevented.

To assess the frequency each cone-opponent group, we developed a framework for relating RGC density to S-cone density. Specifically, our calculations estimate the density of a foveal RGC type based on its expected number per S-cone (*N* ; **Equation 5**). The model was constrained by foveal anatomy and relied on two values from the literature: the proportion of foveal cones that are S-cones and the proportion of foveal RGCs that are midget RGCs. **Figure 6B** shows how the model’s output varies with these parameters and demonstrates that S-cone density is the dominant source of variability. Accordingly, we elected to define a range of estimated RGC densities from the upper and lower bounds of published macaque foveal S-cone densities shown in **Figure 6A** (De Monasterio et al., 1985; Martin and Grünert, 1999).

We first validated our calculations. While foveal small bistratified RGCs receive input from multiple S-cones, there is roughly one per S-cone (Calkins et al., 1998). For *N* = 1, the model estimates a density range of 1.31– 2.21%. This prediction compares well with empirical measurements of small bistratified RGC density (Dacey, 1993; **Figure 6C**), indicating the model is reasonably accurate. The S-OFF neurons also fell at the lower end of this range (**Figure 6D**), consistent with the 1:1 correspondence between S-cones and S-OFF midget RGCs (Klug et al., 2003; Patterson et al., 2019a). By contrast, the observed density of cardinal S-ON neurons (3.7%) was more consistent with 2-3 RGCs per S-cone (**Figure 6C**). Other S-ON neurons are present in the fovea are very rare and unlikely to account for this discrepancy (Grünert and Martin, 2021). Instead, the frequency of S-ON neurons could be due to the location of the imaging plane in the foveal GCL. Specifically, the imaging plane may have been biased towards a superficial region of the GCL where small bistratified RGCs are often located (**Figure 6E**; Patterson et al., 2020b).

We next explored the relationship between the density of S-cones and the rarity of L vs. M ± S neurons. We asked what the density of non-cardinal RGCs would be if there were one from each quadrant in **Figure 3B** per S-cone (L-MS, M-LS, MS-L and LS-M). Given an average of 4 RGCs per S-cone (*N* = 4), the model predicted a density range of 5.2–8.8% of all foveal RGCs (**Figure6D**). The L vs. M ± S neurons (6.6%) fell within this range. While it remains to be determined whether the four groups are equally prevalent, this theoretical estimate provides a useful benchmark as it likely reflects an upper limit on the potential density of non-cardinal RGCs.

In **Figure 6D**, we compare the observed densities of cardinal S-ON, cardinal S-OFF and L vs. M ± S neurons to the density ranges predicted for 1-4 RGCs per S-cone. To facilitate assessment of the full dataset, we also include the previously discussed upper bounds on each group’s density obtained by incorporating in the L vs. S, M vs. S, L+S and M+S neurons from **Figure 4C** that could not be classified unambiguously.

The model’s predictions suggest that cardinal S-ON neurons may be overrepresented (**Figure 6C** and **6E**), possibly due to imaging plane’s location. This explana-tion could account for the high numbers of non-cardinal S-ON neurons (L-MS and M-LS) if their somas are located in the same region of the GCL as the ON midget RGCs (**Figure 6E**). However, assessing this possibility is difficult without knowing the underlying cell type(s). Another possibility suggested by the larger numbers of S-OFF neurons with either Lor M-cone input (**Figure 4C**) is that our stimuli may have been less effective at driving L/M-cone input to S-OFF neurons. Distinguishing between these explanations will be an important direction for future study. Taken together, our data are best interpreted as confirming the presence of L vs. M ± S cone-opponent neurons in primate fovea and paving the way for future studies that can provide more precise density estimates.

## Discussion

A complete account of cone opponency in the retinal output is critical for efforts to understand the neural processing of color. Here we confirmed the existence of foveal neurons with both L vs. M cone opponency and S-cone input. While it remains to be determined how downstream neurons use the responses of the L vs. M ± S neurons, our results confirm that cone-opponent signals in the retinal output are more diverse than suggested by standard color vision models.

While anatomical experiments will be necessary to confirm cell type, we predict the L vs. M ± S neurons are subtypes of midget RGCs, which are the only known pri-mate RGC with L vs. M cone opponency. The random L- and M-cone input to L/M-cone center midget RGCs could explain both the L vs. M ± S neurons in **Figure 3B** and the M+S, L+S, L vs. S and M vs. S neurons in **Figure 4C**. Midget RGCs are the most common primate RGC type; however, there is precedent for overlooked cone opponent subtypes: S-OFF midget RGCs were once highly controversial (Klug et al., 2003; Patterson et al., 2019a). The absence of both S-OFF midget and L vs. M ± S RGCs from some prior surveys could be due to their rarity. Our model suggests their low density can be explained by the rarity of S-cones. Eccentricity may also be an important factor. Unlike most prior studies of cone-opponent primate RGCs, our experiments were performed in the fovea. Higher densities of noncardinal RGCs have been reported in the central retina (De Monasterio et al., 1975). By contrast, the small bistratified RGC density increases with eccentricity (**Figure 6C**), which may contribute to their greater representation in prior work.

The underlying circuit mechanism for combining S-cone signals with L vs. M opponency is not known. There is no evidence for S-cone input to L/M-cone midget bipolar cells or S-ON bipolar cell input to L/M-cone midget RGCs (Schein, 2004; Tsukamoto and Omi, 2016; Patterson et al., 2019a), suggesting horizontal or amacrine cells may play a role. While HI horizontal cell feedback is largely responsible for the L/M-cone surround receptive field, feedback from S-cone-preferring HII horizontal cells is also present (Ahnelt and Kolb, 1994; Packer et al., 2010; Pan and Massey, 2023) and the HII horizontal cells’ feed-forward synapses onto bipolar cells remain largely unexplored (Puller et al., 2014). Known S-cone amacrine cells do not contact midget bipolar cells or RGCs (Patterson et al., 2020a); however, the circuitry of most primate amacrine cell types remains unknown. While one outer retinal mechanism could produce the four forms of non-cardinal cone opponency, an inner retinal origin would likely require distinct S-ON and S-OFF circuit mechanisms. ON/OFF asymmetries are present S-cone retinal circuits and at the level of perception (Dacey et al., 2014; Smithson, 2014). Comparing the spatiotemporal response properties of the non-cardinal neurons with S-ON vs. S-OFF input could narrow the search for potential mechanisms and guide connectomic investigations aimed at uncovering the full circuitry.

Our findings are consistent with a growing appreciation for the importance of rarer RGC types (Masland and Martin, 2007; Gollisch and Meister, 2010) and the capacity for retinal circuits to perform computations typically attributed to the cortex (Patterson et al., 2022a). While the neurons tuned to non-cardinal color directions were relatively rare, their density alone does not rule out a potential contribution to color vision as they exceed the density reported for foveal small bistratified RGCs (**Figure 6D**). Our model suggests that even if each S-cone were represented by all four cone-opponent groups in **Figure 3A**, their density would be just 5.3-8.8% of foveal RGCs. In our dataset, the two S-ON groups (LS-M, MS-L) outnumbered the S-OFF groups (M-LS, L-MS). This imbalance could reflect intrinsic differences between the four groups, the imaging plane’s location within the GCL (**Figure 5C**) or the efficacy of our stimulus for L vs. M opponency in S-OFF neurons (**Figure 4B**). Comparable numbers of L vs. M+S and M vs. L+S RGCs have been reported (De Monasterio et al., 1975). In the LGN, lower numbers of the M–LS neurons have been reported (Tailby et al., 2008). In the primary visual cortex where non-cardinal cone opponency is more prevalent, L vs. M+S neurons outnumber M vs. L+S neurons (Conway, 2001; Horwitz, 2020). A larger sample size will be necessary to distinguish intrinsic differences in the relative densities from experimental factors.

The cardinal directions and their cone inputs have become enormously influential for color vision models (**Figure 1D**). A second stage with three independent orthogonal axes was appealing in its simplicity and agreement with the redundancy reduction hypothesis, a popular theory of neural coding at the time (Barlow, 1961). The cardinal directions’ cone inputs are near optimal for capturing the variance present in the L-, Mand S-cone outputs (Buchsbaum and Gottschalk, 1983; Atick et al., 1992, 1993). However, this hypothesis predicts additional forms of cone opponency are unnecessary because they overlap with the cardinal directions and thus carry redundant spectral information.

However, evidence for greater diversity was present even in the initial work establishing the cardinal directions in the form of residual selectivity for intermediate color directions (Krauskopf et al., 1982) and in the LGN neurons with both L vs. M opponency and S-cone input (Derrington et al., 1984). A reanalysis of the original psychophysical data revealed evidence for additional “higher-order mechanisms” tuned to intermediate color directions (Krauskopf et al., 1986). Subsequent psychophysics provided additional support for higherorder mechanisms and questioned the independence of the cardinal mechanisms (Krauskopf and Farell, 1990; Webster and Mollon, 1991, 1994; Zaidi and Halevy, 1993; Li and Lennie, 1997; Danilova and Mollon, 2012b, 2012a; Emery et al., 2017). As the name “higher-order mechanism” suggests, non-cardinal color tuning is assumed to arise in the cortex (De Valois and De Valois, 1993; Zaidi and Shapiro, 1993). However, our results indicate that some non-cardinal cone opponency is established in the retina and transmitted to the brain in parallel with the outputs of cardinal RGCs.

Despite the consensus that the cardinal directions cannot account for color appearance, the salience of L vs. M and S vs. L+M stimuli at threshold has strong support [reviewed in (Eskew, 2009; Stockman and Brainard, 2010)]. Consistent with previous physiological surveys (De Monasterio and Gouras, 1975; Derrington et al., 1984; Sun et al., 2006; Tailby et al., 2008), most neurons in **Figure 3B** fell along the cardinal axes. Their numerical dominance could provide a simple explanation for the superior detection of L vs. M and S vs. L+M stimuli. While L vs. M mechanisms with S-cone input have been reported in threshold experiments measuring color detection (Stromeyer et al., 1998; Shepard et al., 2017), the strongest psychophysical support for non-cardinal mechanisms comes from suprathreshold experiments measuring color discrimination.

While psychophysics supports macaques as a model of human color vision (De Valois et al., 1974; Stoughton et al., 2012; Gagin et al., 2014; Gelfand and Horwitz, 2018), the possibility of species differences was raised by a recent comparative anatomy study (Kim et al., 2023). Importantly, the recent discussion on species differences emphasizes how the most common macaque RGCs differ from human color appearance, but overlooks their similarities with human color detection. Both are important aspects of color vision requiring explanation at the level of the early visual system. Given the greater diversity of cortical color-tuned neurons (Solomon and Lennie, 2005; Chang et al., 2017; Nigam et al., 2021; Li et al., 2022), retinal circuitry likely provides the beginning of an explanation rather than the final answer.

The L vs. M ± S neurons open new avenues of research for understanding the neural circuits contributing to color perception. Understanding the underlying circuitry and the responses along other stimulus dimensions will be important directions for future research as cone-isolating stimuli alone do not provide a complete basis for developing hypotheses on a cone-opponent RGCs’ role in vision (Patterson et al., 2019b). A comprehensive account should assess how spectral tuning is influenced by spatiotemporal stimulus features and complex, naturalistic stimuli that engage underlying nonlinearities masked by standard linear stimuli.

Despite centuries of study, the underlying mechanisms of color vision remain an unsolved mystery and even foundational hypotheses are debated (Pugh and Kirk, 1986; Gegenfurtner and Hawken, 1996; Knoblauch and Shevell, 2001; Valberg, 2001; Kamar et al., 2019; Conway et al., 2023). The confirmation of L vs. M ± S neurons indicates that models assuming cone opponency in the retinal output is limited to two cardinal mechanisms may require reconsideration. Models where the second stage includes L vs. M ± S neurons can account for many aspects of color perception (Schmidt et al., 2014, 2016; Rezeanu et al., 2023). Furthermore, models exploring the interaction between cardinal and non-cardinal neurons remain largely unexplored. Finally, while color perception is often the default hypothesis for cone-opponent primate RGCs, other visual functions benefit from cone-opponency (Spitschan et al., 2017; Patterson et al., 2022b). Ultimately, a comprehensive account of the neural mechanisms of color perception will require consideration of all cone-opponent RGCs and a wide range of visual functions beyond color appearance.

## Acknowledgements

We thank Amber Walker for animal anesthesia during imaging, David Brainard and Jay Neitz for valuable discussions, the vector core at the Perelman School of Medicine of the University of Pennsylvania, the Genetically-Encoded Neuronal Indicator and Effector (GENIE) project, and the Janelia Research Campus of the Howard Hughes Medical Institute. This work was supported by NIH grants F32-EY032318 (SSP), K99-EY035323 (SSP), P30-EY00131944, R01-EY021166 (WHM), R01-EY031467 (DRW), T32-EY007125 (TG, SSP) and U01-EY025467, Air Force Office of Scientific Research grants FA-9550-22-1-0044 (DRW) and FA-9550-22-1-0167 (DRW), and Research to Prevent Blindness through an unrestricted grant to the Flaum Eye Institute.

## Declaration of Interests

DRW has patents with the University of Rochester (#6199986, #6264328, and #6338559) and has consulted for Warby Parker. QY has patents with the University of Rochester, Canon Inc., and the University of Montana (#9226656, #9406133, and #9454084) and has consulted for Oculus VR and Boston Micromachine Corporation. SSP has a patent application filed with the University of Washington (#17/612061).

### Author Contributions

TG, JEM, WHM, DRW and SSP conceived of the project and designed experiments; TG and JEM collected the data; TG, KK, JEM and SSP analyzed the calcium imaging data; TG developed the crosstalk model; SSP developed the density model; TG and SSP created the figures; KP and QY provided software and hardware; TG and SSP wrote the paper with input from DRW.

